# Potato genotypes differentially alter the expression of *Phytophthora infestans* effectors during PAMP-mediated resistance induction

**DOI:** 10.1101/547984

**Authors:** Cécile Thomas, Pauline Le Boulch, Didier Andrivon, Florence Val

**Author notes:** Author for correspondence: Florence VAL.

## Abstract

- Pathogen recognition by plants *via* pathogen-associated molecular patterns leads to PAMP-triggered immunity. However, pathogens can modulate it *via* the secretion of effectors. We hypothesize that in potato, induced defense triggered by a *Phytophthora infestans* concentrated culture filtrate (CCF) could alter both effector expression and disease severity.
- CCF was sprayed onto three potato genotypes with different resistance levels, before inoculation with *P. infestans*. Symptoms were scored visually at 1-4 dpi, while the expression of defense and effector genes was assessed by qRT-PCR.
- CCF induced most defense genes in Désirée (*PRs, EIN3*) and Bintje (*PRs, PAL* and *POX*), but repressed most defense genes in Rosafolia. On the contrary, CCF induced most effector genes in Rosafolia (*Pi03192, Avrblb2, Avr3a, EPIC2B* and *SNE1*). *INF1* was over-expressed in Bintje, despite its earlier expression in both Désirée and Rosafolia compared to unsprayed controls. *Pi03192* was repressed in Désirée, and was expressed earlier in Rosafolia than in controls. However, induced defense responses by CCF significantly reduced lesion areas at 3 dpi only in Désirée.
- The effectiveness of induced defense thus depends on host genotypes. It results from differential interactions and kinetics of defense and effector genes expressions.

## Introduction

Fungal and oomycete plant pathogens are still responsible for a major fraction of yield losses, despite the use of pesticides which have a negative impact the ecological footprint. Therefore, the current challenge is to find economically viable and environmentally safe methods of disease control (Gururani and Park, 2012). A possible strategy is to exploit plant immunity to enhance plant resistance, combined with quantitative host resistance for more sustainable protection (Walters *et al.*, 2008). For this, it is necessary to better understand the interaction between plant and pathogens. The interactions between plants and pathogens are described as an arms race in the zig-zag model of co-evolution (Jones and Dangl, 2006). Indeed, plants are able to recognize elicitors from pathogens, called pathogen-associated molecular patterns (PAMPs), *via* specific pattern recognition receptors (PRRs). This recognition leads to the induction of general defense responses *via* PAMP-triggered immunity (PTI) in plants. Thus, defense genes induction may result in the accumulation of antimicrobial compounds in the cytosol, as well as in other mechanisms able to enhance plant resistance (Boller and Felix, 2009).

Defense induction by exogenous elicitors ahead of pathogen colonization provides sustainable resistance for plants to a wide range of pathogens (Burketová *et al.*, 2015). Indeed, induced resistance by benzothiadiazole (BTH) protects grapevine from the biotrophic pathogens *Plasmopara viticola* and *Erysiphe necator* (Dufour *et al.*, 2013) and also against the necrotrophic pathogen *Botrytis cinerea* in grapes (Bellée *et al.*, 2018). In potato, β-aminobutyric acid (BABA) reduces *Phytophthora infestans* lesion size and sporangial production on potato leaves (Bengtsson *et al.*, 2014a). Moreover, it protects potato from late blight infection in greenhouse experiments and under field conditions (Liljeroth *et al.*, 2010).

We previously demonstrated that a Concentrated Culture Filtrate (CCF) of *P. infestans*, containing at least four PAMPs (three elicitins and a polysaccharide; Saubeau *et al.*, 2014), induces potato defense genes, and that this induction is genotype-dependent (Saubeau *et al.*, 2016). More recently, Thomas *et al.* (2019) proved that CCF pre-treatment effectiveness depends on *P. infestans* growth rate, and only reduces the lesion in fast-growing strains. Indeed, some pathogens have evolved the ability to modulate or circumvent host cell defenses by secreting apoplastic or cytoplasmic effector proteins, allowing host colonization *via* Effector-triggered susceptibility (ETS; Kamoun, 2006; Dodds and Rathjen, 2010). *P. infestans* has a high rate of effector innovation, thanks to the RXLR gene families within the genome. They are localized in gene-sparse, transposon-rich regions that generate recombinations and diversity (Haas *et al.*, 2009; Raffaele *et al.*, 2010).

In this context, we assumed that potato induced defenses triggered by CCF before infection by *P. infestans* could affect both disease symptoms and effector genes expression. In order to test this hypothesis, we simultaneously assessed in three potato genotypes, pre-treated or not with CCF, (i) the induction of defense genes after inoculation with *P. infestans*, (ii) *P. infestans* effectors involved in counter-defense and (iii) disease symptoms.

## Materials and Methods

### Inoculum and elicitor preparation

*P. infestans* strain—14.P29.03.R—belonging to the 13_A2 clonal lineage was used in the experiments. This strain was characterized as fast-growing by Thomas *et al.* (2019).

*P. infestans* was cultivated for 3 weeks on pea agar, then, sporangia were collected in sterile water. Detached leaflets of cv. Bintje were inoculated with *P. infestans* sporangia to increase its pathogenicity. Six days after incubation of infected leaflets in humid chamber, sporangia were collected in sterile water, and inoculum was prepared and adjusted to a final concentration of 5.10^4^ sporangia.mL^−1^ as described previously by Montarry *et al.* (2007), adapted by Thomas *et al.* (2019).

*P. infestans* concentrated culture filtrate (CCF) was prepared as described by Desender *et al.* (2006) from strain 14.P29.03.R, filtrated on sterile gauze and lyophilized for 72 hours. Then, it was diluted in water at 8 mg.mL^−1^ and 0.1% Tween20 (Saubeau *et al.*, 2016).

### Cultivation of plants

Tubers of *Solanum tuberosum* (L.) genotypes Bintje, Désirée and Rosafolia, chosen according to their various level of susceptibility to *P. infestans* (Clément, 2011), were grown in a greenhouse as described by Thomas *et al.* (2019). Plants were watered once a week with a nutrient solution (NPK 15/10/15; Hakaphos).

### Experimental design

Four weeks following planting, two independent experiments (only one for Rosafolia) were carried out for each genotype, with the same experimental design. Eighteen plants were sprayed to runoff with either CCF at 8 mg.ml^−1^ (+0.1% Tween20) or sterile water (+0.1% Tween20, control). Each experiment tested eight conditions for each genotype, *i.e*. all possible combinations of two leaflet pre-treatments (water or CCF) inoculated with *P. infestans*, and four observation dates (one, two, three and four days post inoculation). Each treatment involved 12 leaflets.

### Sampling of leaflets and inoculation

Forty-eight hours (48 h) after pre-treatment, leaflets were picked at random from the fifth and sixth leaf levels below the apex. Sampling of leaflets and inoculation with 20 μL of *P. infestans* sporangial suspensions at 5.10^4^ sporangia.mL^−1^ were done as described by Thomas *et al.* (2019).

### Assessment of potato defense genes and *Phytophthora infestans* effector genes by high throughput quantitative reverse transcription-polymerase chain reaction (qRT-PCR)

The expression of fourteen defense genes (**Table 1**) and of nine effector genes (**Table 2**) was assessed at 1, 2, 3 and 4 days post inoculation (dpi). The 12 leaflets from each condition were sampled as described by Thomas *et al.* (2019), lyophilized 24 h and ground at room temperature to a fine powder with Fastprep. This lyophilized powder was distributed as 15 mg aliquots into 2 mL cryogenic tubes. RNA was then extracted independently from two of these aliquots), as described by Saubeau *et al.* (2016). The integrity of the RNA was evaluated by electrophoresis on 1% agarose gel before conversion into cDNA.

**Table 1.**
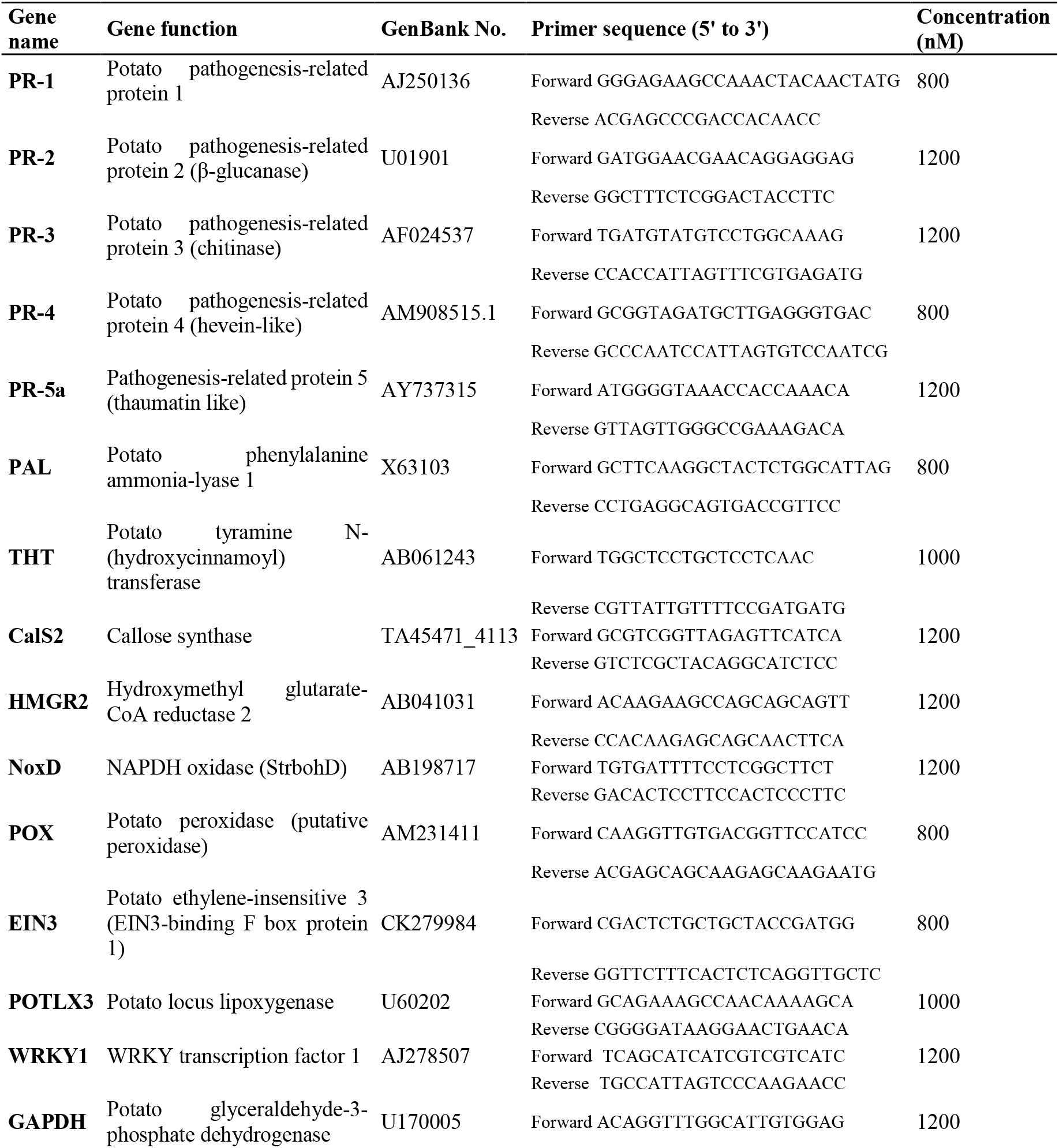
Selected defense genes and corresponding primer sets used for analysis of transcript profiles from *Solanum tuberosum* leaves inoculated with *P. infestans* (Saubeau, 2014)

**Table 2.**
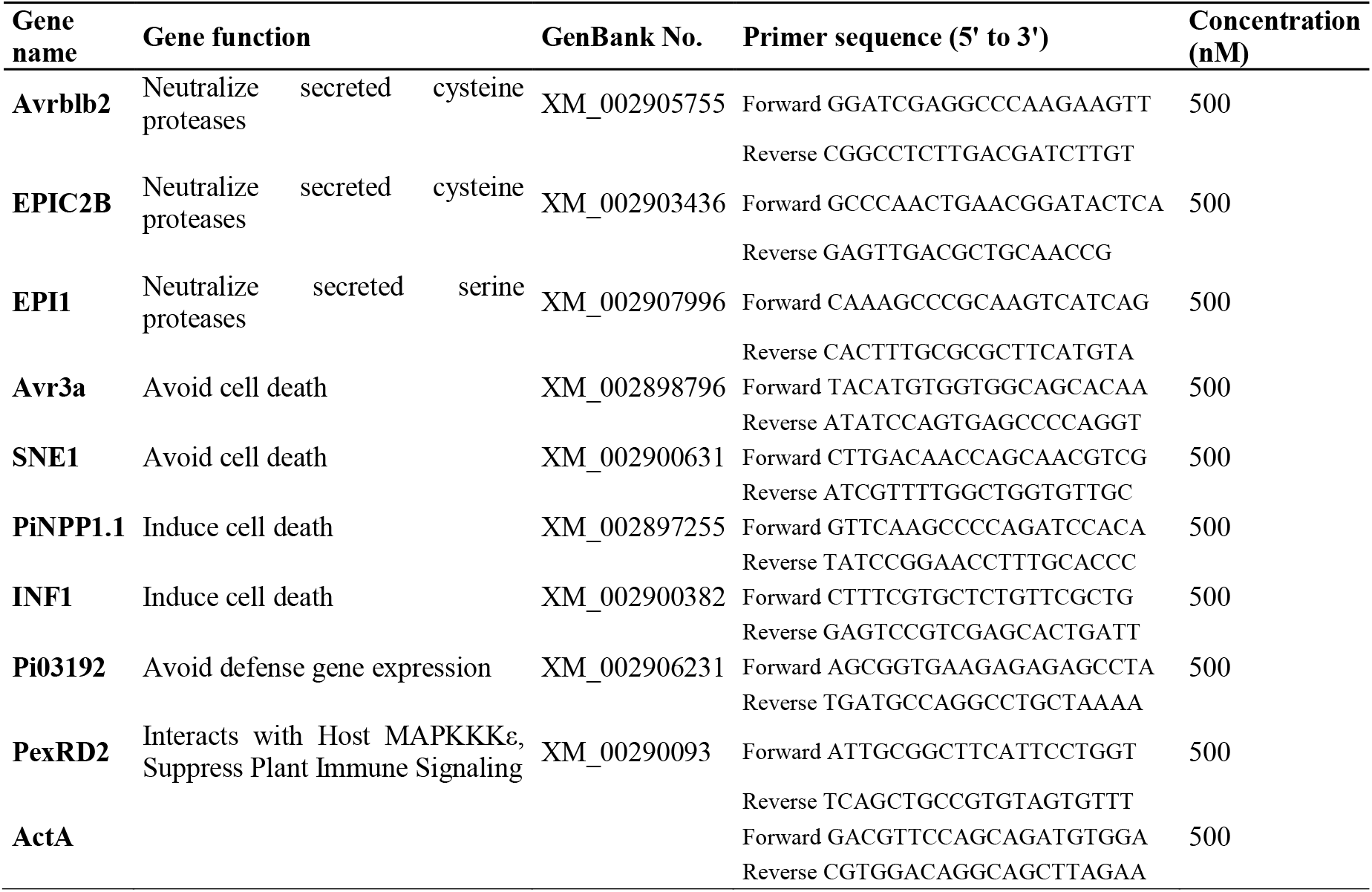
Selected effector genes and corresponding primer sets used for analysis of transcript profiles from *Solanum tuberosum* leaves inoculated with *P. infestans* (A. Kröner and R. Mabon, personal communication)

Each cDNA sample was then amplified in four qRT-PCR reactions for each defense and effector gene targeted (qPCR replicates). The expression of fourteen potato defense genes (**Table 1**) and of nine *P. infestans* effector genes (**Table 2**) was simultaneously analyzed on the same samples at 1, 2, 3 and 4 dpi with qRT-PCR in leaf samples sprayed with CCF or water and inoculated with *P. infestans*.

High-throughput qRT-PCR reactions were performed using the WaferGen SmartChip Real-time PCR system (5184-nanowells chip, WaferGen Bio-systems and Takara Bio Inc. USA). qRT-PCR reaction volumes of 100 nL consisted of 1 X LightCycler 480 SYBR Green I Master (Roche Diagnostics France), a cDNA sample of 1 ng.μL^−1^, 500-1200 nM primers mix (desalt purification, all provided by Sigma-Aldrich Chimie S.a.r.l France, **Tables 1 and 2**). qPCR mixtures were dispensed by SmartChip Multisample Nanodispenser (MSND) into two SmartChip patterns to assess respectively defense transcripts (36 assays × 144 samples) and effector transcripts (24 assays × 216 samples). For each experiment and treatment, cDNA were taken from two aliquots into four qPCR reactions.

Amplifications were performed in SmartChip Cycler under the following cycling conditions: 5 min at 95°C; 50 cycles of 5 s at 95°C, 30 s at 60°C and 30 s at 72 °C for defense genes; 15 min at 95°C; 50 cycles of 15 s at 95°C, 30 s at 62°C and 30 s at 72 °C for effector genes. Melting curve analyses were performed at the end of qPCR for both defense and effector genes from 62°C to 95°C.

qRT-PCR results were analyzed using SmartChip qPCR Software (v. 2.8.6.1) for melting and amplification curves. Wells with multiple melting peak as well as those with bad amplification curves were discarded. Then, transcript accumulation of defense and effector genes was calculated by using the relative quantification method described by Pfaffl (2001). To calculate qRT-PCR efficiencies from calibration curves (*E*= 10^[–1/slope]^) and thus perform the inter-chip normalization, each qRT-PCR run x primer set included a dilution range. Genes with amplification effectiveness beyond the range 1.7 − 2.2 were discarded (*i.e.* defense genes *THT, CalS2, WRKY1* and effector genes *EPI1, PiNPP1.1, PexRD2*). Expressions of the targeted genes were then normalized with the reference gene *GAPDH* (defense) or *ActA* (effectors). Normalized transcript accumulation data were expressed for each condition from duplicates of two independent experiments.

### Tracking pathogen development by measuring lesion area

Pathogen development was evaluated at 2, 3 and 4 dpi by measuring lesion on the abaxial surface of infected leaflets pretreated or not with CCF with a caliper. Lesion area was, then, calculated with the formula used by Vleeshouwers *et al.* (2000), assuming an elliptic shape: Lesion area = length × width × π × ¼.

### Statistical analyses

Statistical analyses were performed using the statistical software R version 3.5.1. Potato genotype and CCF pre-treatment effects were tested at each point of kinetic on lesion area (symptoms) with one-way variance analyses (ANOVA, function ‘aov’). Residuals normality and variances homogeneity were verified. These effects were also tested at each point of kinetic on transcript accumulation (plant defense genes and *P. infestans* effector genes expression) with Kruskal-Wallis analyses. Null hypotheses were rejected if *P value* < 0.1.

## Results

### Phytophthora infestans alone induces defense responses depending on the genotype

The global analysis of induced defenses by *P. infestans* showed a difference in expression between genotypes (**Fig. 1**). Indeed, Désirée was characterized by the expression of *PR-2, PR- 4, PR-5a* and *POX*, whereas Bintje was characterized by the expression of *PR-3, EIN3, POTLX3* and *NoxD*. Furthermore, defense gene expression was generally lower in Rosafolia, except for *PR-1*.

**Fig. 1.**
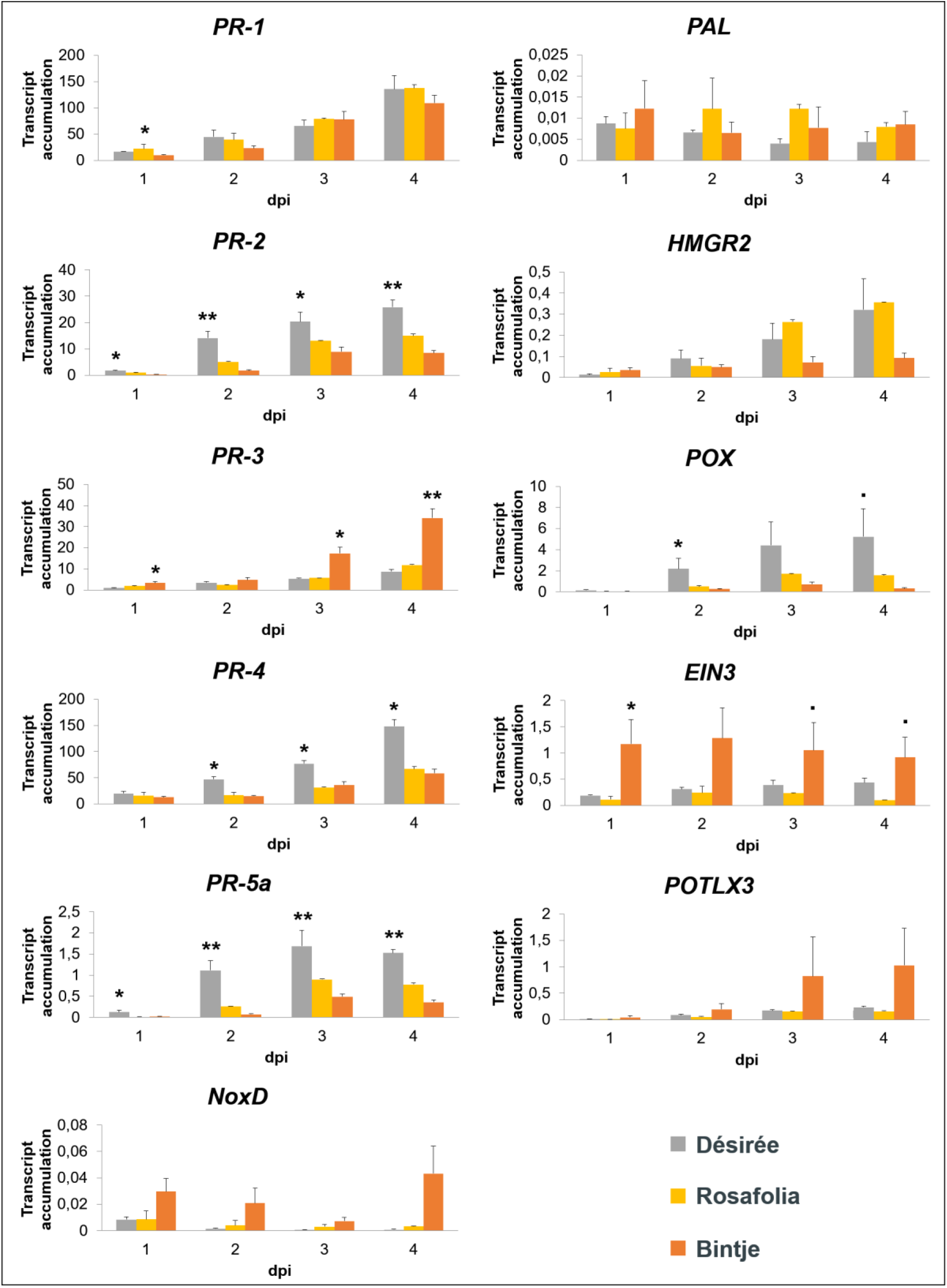
Genotype effect on potato defense genes expression after inoculation with *Phytophthora infestans*. Transcript accumulation of defense genes involved in various signaling pathways in potato leaflets Désirée, Rosafolia and Bintje sprayed with water (control), from 1 to 4 days post inoculation (dpi) with *P. infestans*. Data represent the mean + standard error of the mean (SEM), n=2; results were obtained in two independent experiments (only one experiment for Rosafolia). The genotype effect was tested for each day (Kruskal Wallis: • *P value* ≤ 0.1, * *P value* ≤ 0.05, ** *P value* ≤ 0.01).

There was a higher accumulation of *PR-2, PR-4, PR-5a* and *POX* in Désirée than in Bintje and Rosafolia. It was significant for *PR-2* and *PR-5a* from 1 to 4 dpi (*P value* ≤ 0.05 and *P value* ≤ 0.04; while it was significant for *PR-4* and *POX* from 2 to 4 dpi (*P value* 0.03 and *P value* ≤ 0.06). On the contrary, there was a significantly higher accumulation of *PR-3* and *EIN3* in Bintje than in Désirée and Rosafolia at 1, 3 and 4 dpi (*P value* ≤ 0.03 and *P value* ≤ 0.10) and of *NoxD* from 1 to 3 dpi (*P value* ≤ 0.13). On the other hand, there was a higher accumulation of *PR-1* in Rosafolia than in Désirée and Bintje at 1 dpi (*P value* 0.04).

### CCF pre-treatment modulates the expression of some defense genes depending on the genotype

The global analysis of induced defenses by CCF pre-treatment showed a difference in expression between genotypes (**Fig. 2**). Indeed, CCF increased the expression of the most targeted defense genes in Bintje (*PR-1, PR-2, PR-3, PR-4, PR-5a, PAL* and *POX*) and in Désirée (*PR-1, PR-2, PR-3, PR-4, and EIN*3). On the contrary, it decreased the expression of the most targeted genes (*PR-2, PR-5a, PAL, POX, HMGR2* and *POTLX3*) in Rosafolia. However, CCF induced the expression of *PR-1, PR-3* and *PR-4* in Rosafolia.

**Fig. 2.**
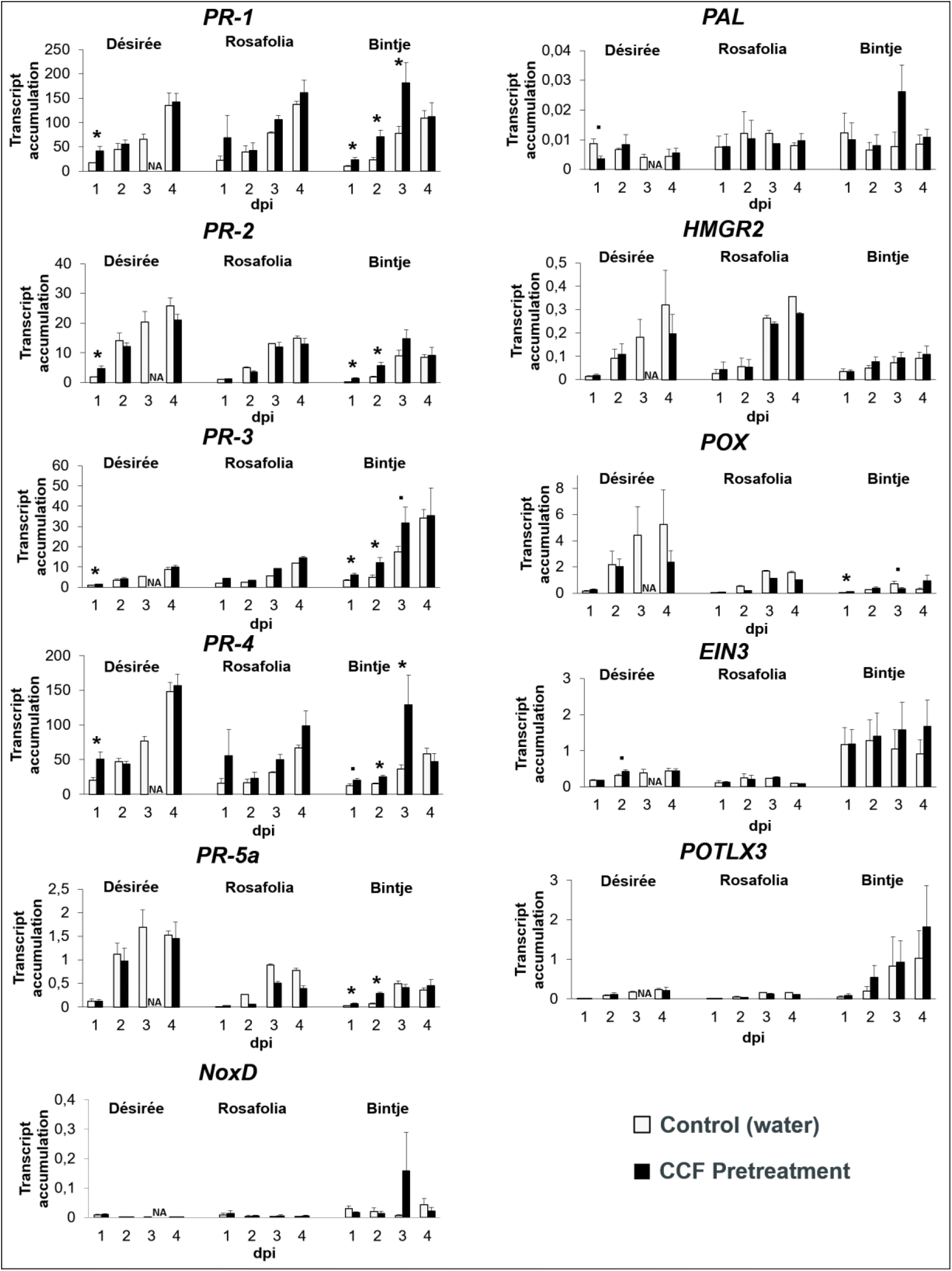
CCF pre-treatment effect on potato defense genes expression after inoculation with *Phytophthora infestans*. Transcript accumulation of defense genes involved in various signaling pathways in potato leaflets Désirée, Rosafolia and Bintje sprayed with either CCF or water (control), from 1 to 4 days post inoculation (dpi) with *P. infestans*. Data represent the mean + standard error of the mean (SEM), n=2; results were obtained in two independent experiments (only one experiment for Rosafolia). The CCF pre-treatment effect was tested for each day, and each genotype (Kruskal Wallis: • *P value* ≤ 0.1, * *P value* ≤ 0.05).

CCF pre-treatment increased significantly the accumulation of most *PRs* in both Désirée and Bintje, but affected differently according to the amount of time. Indeed, *PR-1, PR-2, PR-3* and *PR-4* were more accumulated at 1 dpi in Désirée (*P value* 0.03, but this increase was maintained longer in Bintje, from 1 to 2 dpi for *PR-2* and *PR-5a* (*P value* ≤ 0.04), and from 1 to 3 dpi for *PR-1, PR-3* and *PR-4* (*P value* ≤ 0.08). In Rosafolia, the expression of *PR-3*, and of *PR-4* increased respectively from 1 to 4 dpi, and from 3 and 4 dpi, but it was transient for *PR-1* at 3 dpi, and for *PR-5a* at 1 dpi (*P value* 0.12). On the contrary, CCF decreased the expression of *PR-2* at 2 dpi and *PR-5a* from 2 to 4 dpi in this same genotype (*P value* 0.12). CCF significantly reduced *PAL* accumulation at 1 dpi in Désirée (*P value* 0.08) and at 3 dpi in Rosafolia (*P value* 0.12), whereas *PAL* accumulation increased at 3 dpi in Bintje (*P value* 0.14). There was a significant increase of *POX* accumulation at 1 dpi in Bintje (*P value* 0.02), whereas there was a significant decrease in accumulation at 3 dpi (*P value* 0.08). Moreover, CCF decreased the expression of *POX* from 2 to 4 dpi in Rosafolia (*P value* 0.12). We also observed a significant increase of *EIN3* expression at 2 dpi in Désirée (*P value* 0.07) and at 3 dpi in Rosafolia (*P value* 0.12), whereas it reduced *EIN3* expression at 4 dpi in Rosafolia (*P value* 0.12). CCF decreased the expression of *HMGR2* and *POTLX3* only in Rosafolia from 3 to 4 dpi (*P value* 0.12).

### Phytophthora infestans induces the expression of effector genes depending on the genotype

The global analysis of effectors induction by *P. infestans* showed a difference in expression between genotypes (**Fig. 3**).

**Fig. 3.**
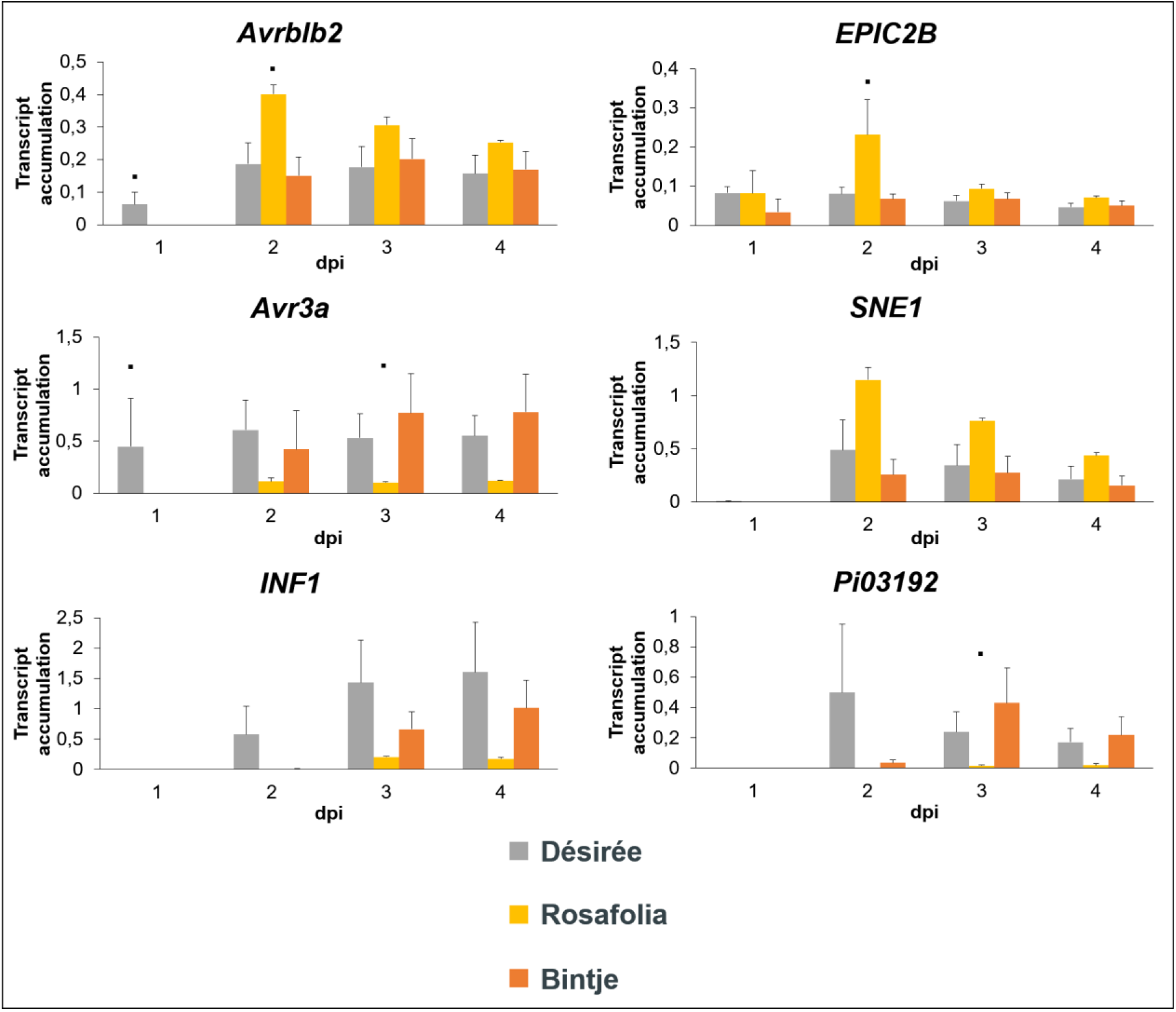
Genotype effect on the expression of *Phytophthora infestans* effectors involved in counter-defense. Transcript accumulation of effector genes with different functions in potato leaflets Désirée, Rosafolia and Bintje sprayed with water (control), from 1 to 4 days post inoculation (dpi) with *P. infestans*. Data represent the mean + standard error of the mean (SEM), n=2; results were obtained in two independent experiments (only one experiment for Rosafolia). The CCF pre-treatment effect was tested for each day, each genotype and each treatment (Kruskal Wallis: • *P value* ≤ 0.1).

After inoculation with *P. infestans, Avrblb2* and *Avr3a* were expressed one day earlier in Désirée (from 1 dpi, *P value* 0.10) than in Rosafolia and Bintje (from 2 dpi). Moreover, *INF1* and *Pi03192* were expressed one day earlier in both Désirée and Bintje (from 2 dpi) than in Rosafolia (from 3 dpi). However, *EPIC2B* and *SNE1* were respectively expressed from 1 and 2 dpi in the three genotypes.

*Avrblb2* and *EPIC2B* were significantly more expressed at 2 dpi in Rosafolia than in Bintje and Désirée (*P value* 0.08). Moreover, *SNE1* was more expressed from 2 to 4 dpi in Rosafolia (*P value* ≤ 0.14). On the contrary, *Avr3a* was less expressed at 3 and 4 dpi in Rosafolia (*P value* ≤ 0.11), and *INF1* was less expressed in this genotype at 4 dpi (*P value* 0.18). On the other hand, *INF1* and *Pi03192* were less expressed at 2 dpi in Bintje than in Désirée (*P value* 0.14 and 0.11), but *Pi03192* was significantly less expressed at 3 dpi in Rosafolia than in Bintje and Désirée (*P value* 0.10).

### CCF pre-treatment modulates the expression of effector genes depending on the genotype and on the amount of time (Fig. 4)

*Pi03192* was expressed one day earlier in Rosafolia after CCF pre-treatment (from 2 dpi rather than from 3 dpi) and was over-expressed at 3 dpi in this genotype (*P value* 0.12). In contrast, CCF did not alter the kinetic of expression of *Pi03192* in Bintje and Désirée, while it was repressed at 2 dpi in Désirée (*P value* 0.02). *EPIC2B* was expressed one day later in Rosafolia (from 2 dpi rather than from 1 dpi) after CCF pre-treatment but was over-expressed at 4 dpi (*P value* 0.12). In contrast, CCF did not alter the kinetic or the level of expression of *EPIC2B* in neither Bintje nor Désirée. *INF1* was expressed one day earlier in both Désirée (1 dpi) and Rosafolia (2 dpi) after CCF pre-treatment, while it was over-expressed at 2 dpi in only Bintje (*P value* 0.02). However, CCF repressed *INF1* at 3 dpi in only Rosafolia (*P value* 0.12).

**Fig. 4.**
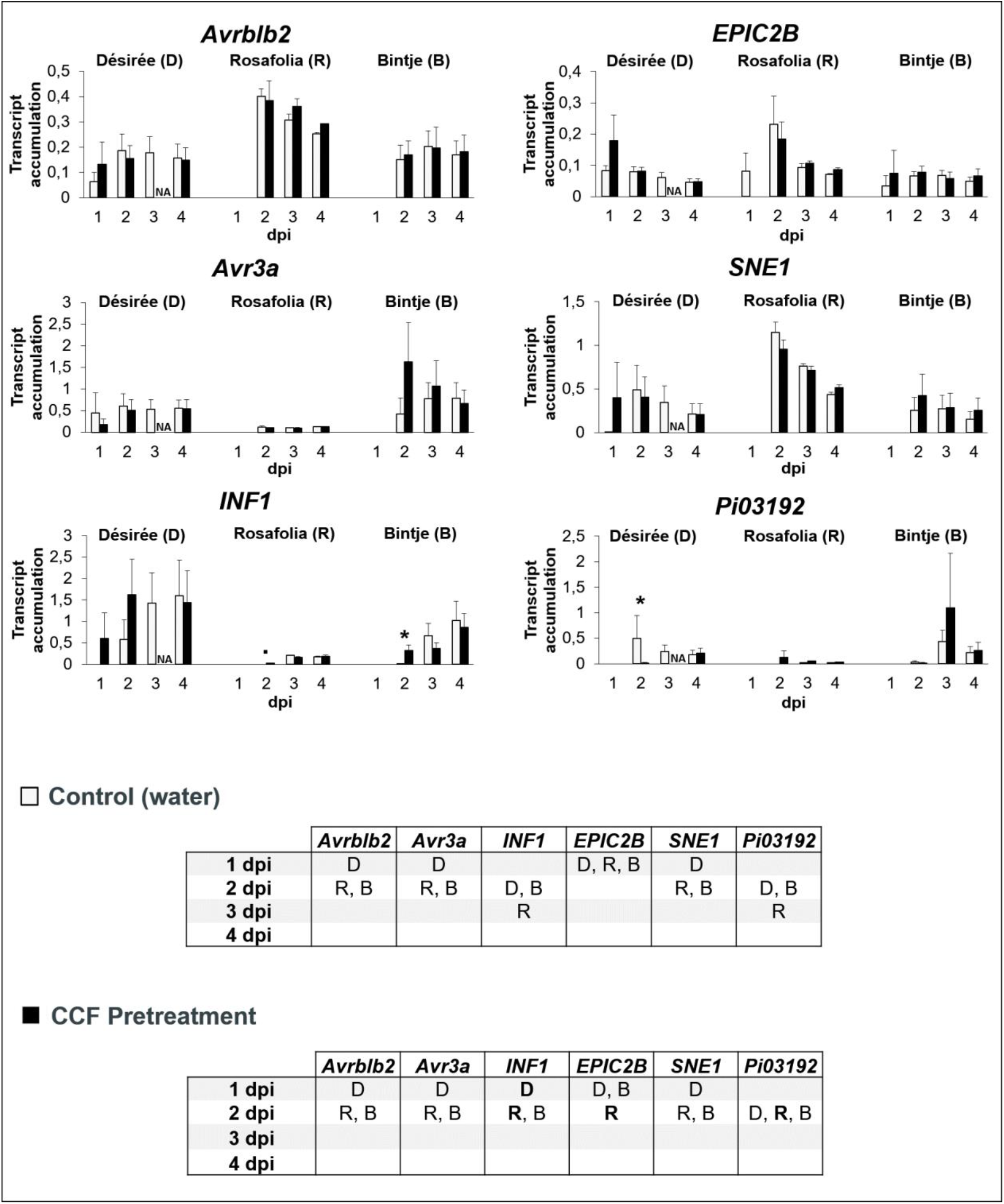
CCF pre-treatment effect on the expression of *Phytophthora infestans* effectors involved in counter-defense. Transcript accumulation of effector genes with different functions in potato leaflets Désirée (D), Rosafolia (R) and Bintje (B) sprayed with either CCF or water (control), from 1 to 4 days post inoculation (dpi) with *P. infestans*. Data represent the mean + standard error of the mean (SEM), n=2; results were obtained in two independent experiments (only one experiment for Rosafolia). The CCF pre-treatment effect was tested for each day, and each genotype (Kruskal Wallis: • *P value* ≤ 0.1, * *P value* ≤ 0.05). The table summarizes the first day of effectors expression, for each genotype with either CCF or water (control).

On the other hand, the kinetic and the level of expression for *Avrblb2, Avr3a* and *SNE1* were not altered by CCF pre-treatment. However, *Avrblb2* was over-expressed from 3 to 4 dpi in Rosafolia after CCF pre-treatment, while *Avr3a* and *SNE1* were over-expressed at 4 dpi (*P value* 0.12) in this genotype.

### The effectiveness of induced defenses depends on the genotype

Without pre-treatment, lesion area was larger in Désirée than in the other genotypes from 3 to 4 dpi (*P value* ≤ 2.10^−6^). However, CCF pre-treatment prior to inoculation with *P. infestans* significantly reduced lesion area at 3 dpi only in Désirée (*P value* 0.02, **Fig. 5 A, B**). There was only a tendency at 3 dpi in Rosafolia (*P value* 0.26, **Fig. 5 A, B**) and at 4 dpi in both Désirée and Bintje (*P value* 0.18 and 0.27, **Fig. 5 A, C**).

**Fig. 5.**
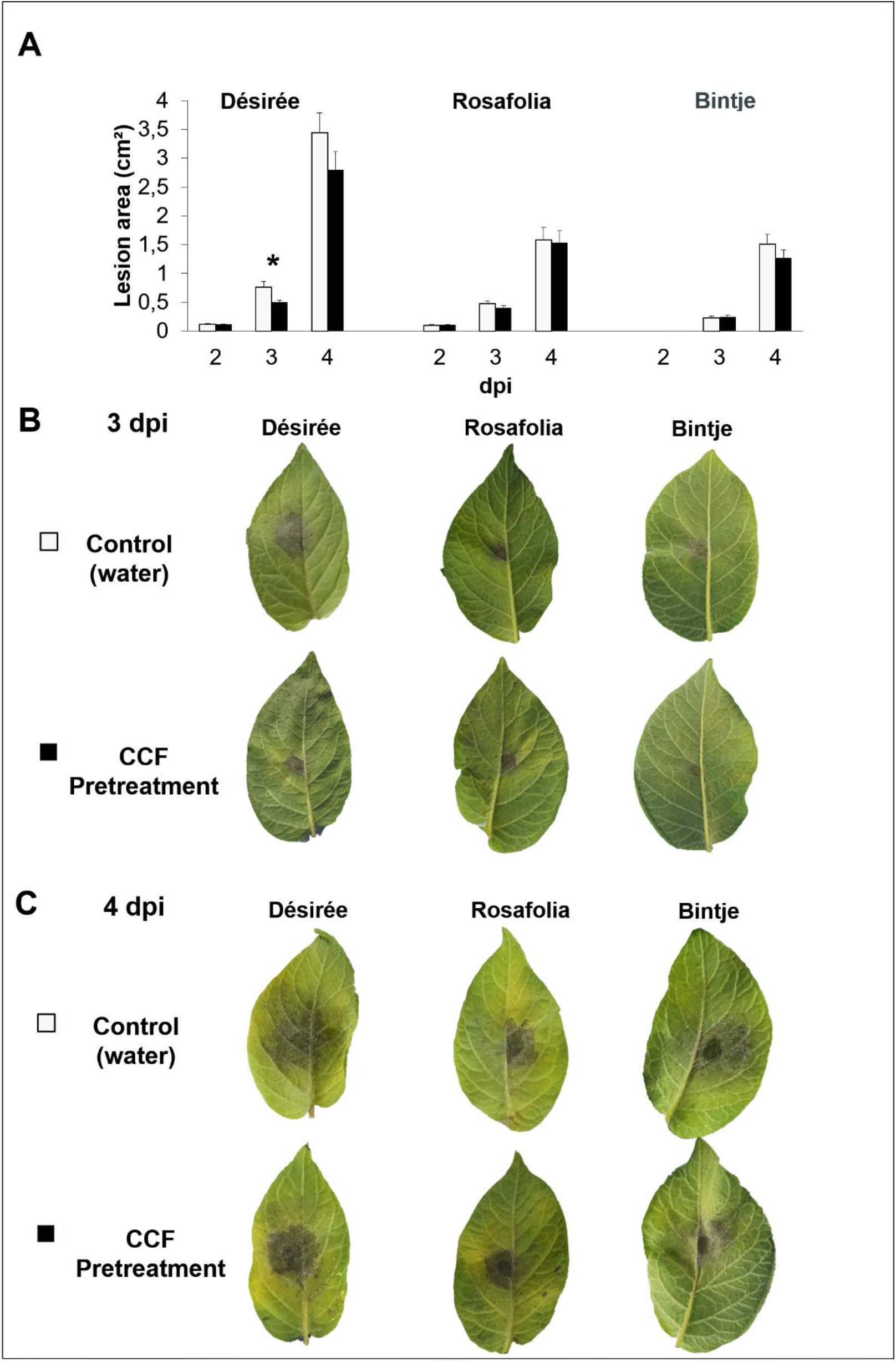
CCF pre-treatment effect on *Phytophthora infestans* symptoms. Lesion area on potato leaflets Désirée, Rosafolia and Bintje sprayed with either CCF or water (control), (**A**) from 2 to 4 days post inoculation (dpi), with *P. infestans* at 3 dpi (**B**) and at 4 dpi (**C**). Data represent the mean + standard error of the mean (SEM), n=12; results were obtained in two independent experiments (only one experiment for Rosafolia). The CCF pre-treatment effect was tested for each day, and each genotype (ANOVA: * *P value* ≤ 0.05).

## Discussion

Our working hypothesis was that the induced defenses triggered in potato treated with CCF before infection by *P. infestans* would affect both disease symptoms and effector genes expressions. The analysis of induced defenses by PAMPs of *P. infestans* indeed showed a difference in expression levels, but also kinetics, between potato genotypes. Indeed, CCF increased the expression of potato defense genes transiently in Désirée, and for a longer period in Bintje. By contrast, it repressed most defense genes in Rosafolia, although transcript accumulation varied over time for this genotype. These results are consistent with those of Saubeau *et al.* (2016), who showed that the expression patterns of defense genes induced by CCF differed according to genotypes rather than to field resistance levels. Interestingly, defense induction by exogenous elicitors as BABA or phosphite was often higher in resistant than in susceptible potato genotypes (Liljeroth *et al.*, 2010; Lim *et al.*, 2013; Bengtsson *et al.*, 2014a).

Our results showed that CCF mainly induced the expression of PR genes, while the expression of other defense genes was consistently lower. Our results are consistent with those obtained with BABA and phosphite. Indeed, BABA induced a major accumulation of *PR-1* transcripts in Bintje and Ovatio (Bengtsson *et al.*, 2014a) and of PR-1 and PR-2 proteins in Désirée (Bengtsson *et al.*, 2014b). Moreover, phosphite induced the accumulation of three secretory proteins such as PR-2, PR-3, and lipase in the susceptible genotype Russet Burbank (Lim *et al.*, 2013). PR proteins are used as markers for pathogen responses and for induction by elicitors, because they play an important role in antimicrobial properties on local infection sites. Indeed, they act as hydrolytic enzymes, contributing directly to the degradation of pathogen cell walls and to the disruption of pathogen membrane integrity (van Loon *et al.*, 2006).

It is noteworthy that *P. infestans* alone induced a higher accumulation of some defense genes in Désirée and of other defense genes in Bintje. However, the expression of defense genes was generally lower in Rosafolia than in the other genotypes. The fact that these three genotypes come from different genetic backgrounds could explain the different levels of defense genes induction after inoculation with *P. infestans*: (i) Désirée descends from the cross between Urgenta and Depesche, (ii) Bintje derives from the cross between Munstersen and Fransen, and Rosafolia derives from the cross between Centifolia and Parnassia (http://www.europotato.org). These three genotypes differ in susceptibility to *P. infestans* (http://www.europotato.org), and in resistance components (Clément *et al.*, 2010). The reduced lesion area in Rosafolia compared to the other two genotypes could be due to constitutive resistance, such as physical - cuticule (Jeffree, 1996) - or chemical barriers - phytoanticipins (Nicholson and Hammerschmidt, 1992). Since Thomas *et al.* (2019) demonstrated that slow-growing strains of *P. infestans* caused a lower defense induction than their fast-growing counterparts (probably due to a lower secretion of PAMPs), this could explain why we observed a lower expression of defense gene in this genotype.

We proved that *P. infestans* infection induced the accumulation of six targeted effectors involved in counter defense, which also showed a difference in expression between genotypes (**Fig. 3**). Indeed, *Avrblb2, EPIC2B* and *SNE1* were more accumulated in Rosafolia than in both Bintje and Désirée, whereas the expression of *Avr3a, INF1* and *Pi03192* was only lower in Rosafolia. The kinetics of *P. infestans* effectors expression was also genotype-dependent, which sustains our initial hypothesis. Indeed, *Avrblb2* and *Avr3a* were expressed one day earlier in Désirée than in the other genotypes, whereas *INF1* and *Pi03192* were expressed one day earlier in both Désirée and Bintje. However, CCF pre-treatment induced an earlier expression of *INF1* in both Désirée and Rosafolia and *Pi03192* only in Rosafolia. *EPIC2B* was expressed later in this pre-treated genotype compared to *P. infestans* alone. Moreover, CCF induced most effector genes over-expression in Rosafolia (*Pi03192, Avrblb2, Avr3a, EPIC2B* and *SNE1*) whereas *INF1* was only over-expressed in Bintje and *Pi03192* was only repressed in Désirée.

*EPIC2B* and *Avrblb2* could be responsible for the lower expression of *PR* genes in Rosafolia, compared to Bintje and Désirée after inoculation with *P. infestans*. Indeed, Tian *et al.* (2007) and Bozkurt *et al.* (2011) showed that in the pathosystem *P. infestans*-tomato, *EPIC2B* and *Avrblb2* are involved in counter-defense against apoplast-localized host proteases (respectively C14 and PIP1), promoting *P. infestans* virulence. Moreover, PIP1 is regulated with a large number of SA-induced PR-proteins such as PR-1 and PR-2 (Zhao *et al.*, 2003). These effectors were over-expressed after CCF pretreatment, and thus could explain the repression of *PR-2* and *PR-5a* in Rosafolia according to Rose *et al.* (2002) who showed that a glucanase inhibitor protein of *P. sojae* inhibits a soybean β-1,3-glucanase belonging to the PR-2 class. The over-expression of *Pi03192* could be a possible explanation for the repression of most defense genes. This effector manipulates host gene expression by downregulating defense related genes (McLellan *et al.*, 2013). In the same genotype, the higher expression of *SNE1*, a suppressor of programmed cell death involved in biotrophy (Kelley *et al.*, 2010), could be responsible for the smaller and less necrotic lesions. This result is also consistent with the lower expression of *INF1*, a necrosis-inducting effector (Kamoun *et al.*, 1998).

The interaction between induced defenses in potato and the modulation of the expression of *P. infestans* effectors by CCF pre-treatment could result in different effectiveness depending on the genotype. The induction of defense genes by CCF before inoculation with *P. infestans* only led to a significant reduction of lesion area at 3 dpi on Désirée. This result is consistent with the behavior of this genotype in field trial, because the treatment of Désirée with various defense inducers has led to a reduction in symptoms. Moreover, it is interesting to note that in Bintje, unlike in Desiree, the induction of defense genes did not lead to a significant reduction of lesion area, meaning that defenses induced by CCF-pre-treatment are not effective for all genotypes. However, as described by Moushib *et al.* (2013) and Bengtsson *et al.* (2014a), pre-treatment with sugar beet extract or BABA reduces lesion area in all genotypes, regardless of their resistance level (Désirée, Bintje, Ovatio). In Rosafolia, CCF pre-treatment led to the over-expression of five effector genes, but repressed most defense genes. This could explain the lack of effectiveness of CCF in this genotype.

This study therefore largely confirmed our initial hypothesis, by showing that the differential interplay between host response to elicitation and infection in the expression of defense genes on one side, and of pathogen effector genes on the other side, largely condition the effectiveness of induced defense on symptom development. It further strengthened earlier observations (Saubeau *et al.*, 2016) about the genotype dependence of such expressions and outcomes. Together these results provide a better understanding of the conditions for effective induced resistance in potato, and open new ground for searching potato germplasm for genetic determinants favoring such resistance.

## Acknowledgements

The authors thank Romain Mabon and Alexander Kröner who developped the primers and qRT-PCR protocol for effectors analysis, and Viviane Lecomte for microbiology assays support. We are most grateful to Biogenouest Genomics and the Human & Environmental Genomics core facility of Rennes (Biosit/OSUR) for its technical support. The authors are grateful to the INRA Genetic Resources Center BrACySol of Ploudaniel for providing the plant material used in the tests. C. Thomas is supported by a CIFRE PhD grant from Bretagne Plants Innovation, acting on behalf of ACVNPT (Association des Créateurs de Variétés Nouvelles de Pomme de Terre). The authors also acknowledge N. Seal, English consultant, for the manuscript correction.

## Author Contribution

C.T., F.V and D.A. developed the concept. C.T. designed and planned the experiments. C.T. and P.L.B. carried out the experiments. C.T. analyzed data with contribution from P.L.B. for the analysis of *P. infestans* symptoms. C.T., P.L.B., F.V and D.A. interpreted and discussed the results; C.T. wrote the manuscript with contributions from F.V., D.A and P.L.B.

